# Adaptive landscape of the metallo-β-lactamase NDM to emerging β-lactam-based therapies

**DOI:** 10.64898/2026.07.10.737697

**Authors:** Fernanda D. Romero, Salvador I. Drusin, Guillermo Bahr, Gastón González, Robert A. Bonomo, Diego M. Moreno, Lisandro J. González, Alejandro J. Vila

## Abstract

The New Delhi metallo-β-lactamase (NDM) is a major determinant of carbapenem resistance. This has prompted the development of novel therapeutic strategies to treat NDM producers. Since these therapies impose new selective pressures, their clinical deployment may favor NDM variants carrying escape mutations. To anticipate these events and inform future therapeutic decisions, we explored the evolutionary landscape of NDM-1 under different selective constraints. To this end, we generated a highly diverse library of *bla*_NDM_ variants and challenged it with zinc starvation, antibiotics or β-lactam/β-lactamase inhibitor combinations that represent novel and emerging therapies. Selection under zinc limitation and cefotaxime identified mutational trajectories that recapitulate clinical NDM evolution, validating the diversity of the library and its predictive nature. Drug-specific selections revealed sharply different evolutionary pathways. Mecillinam selected a narrow evolutionary pathway centered on residues N220 and M67 which enhance productive active-site interactions with this penicillin. Cefepime/taniborbactam selected multiple escape routes, dominated by substitutions at E152 and, secondarily, K211, that impair productive interaction with taniborbactam. In contrast, cefiderocol/xeruborbactam and aztreonam/avibactam failed to select NDM variants conferring an improved resistance phenotype. These results show that NDM evolution is constrained by the chemistry of each therapeutic challenge. Substrate adaptation is possible for mecillinam, inhibitor escape is readily accessible for taniborbactam, whereas aztreonam- and xeruborbactam-based strategies impose high evolutionary barriers on NDM. Mapping drug-specific evolutionary landscapes can help anticipate resistance before clinical deployment and prioritize therapeutic strategies less likely to drive NDM-mediated escape.

**Importance:** Carbapenem-resistant infections by Enterobacterales are increasingly difficult to treat because bacteria can destroy some of the most powerful antibiotics used in the clinic. This study focuses on *Escherichia coli* carrying variants of the New Delhi metallo-β-lactamase, an enzyme that enables bacteria to resist carbapenem antibiotics. New therapies are being developed to overcome this problem, but bacteria may evolve again when exposed to these treatments. Here, we tested how New Delhi metallo-β-lactamase can adapt under the evolutionary pressure of last-resort therapies. The results show that not all treatments carry the same evolutionary risk. Some drugs allow the enzyme to adapt through specific mutations, whereas other drug combinations make such escape much harder. This work helps predict which treatments are more likely to remain effective and which may more readily select resistance. This information can guide the design and use of future therapies against resistant bacterial infections.

## Introduction

Carbapenem-resistant Gram-negative infections represent one of the most difficult challenges to contemporary antimicrobial therapy (1,2). A major driver of this problem is the dissemination of genes coding for metallo-β-lactamases (MBLs), zinc-dependent enzymes that hydrolyze most β-lactams, including carbapenems (3–7). Currently, MBLs are not inhibited by clinically approved β-lactamase inhibitors that target serine β-lactamases (SBLs) (5,8,9). In particular, the New Delhi metallo-β-lactamase (NDM) is becoming more prevalent worldwide since the *bla*_NDM_ gene is present on mobile genetic elements that include other resistance determinants (5,10–12). NDM also possesses unique protein features such as a broad resistance phenotype and its cellular localization, being a membrane-anchored protein (13). This cellular localization favors the stability of NDM in the periplasm, does not induce any fitness cost in several bacterial hosts, and favors its secretion into outer membrane vesicles (14,15). The synergy of these genetic and protein traits clearly favors the dissemination of NDM among different bacterial pathogens. Fortunately, recent developments offer new therapeutic possibilities for infections caused by MBL-producing bacteria.

Aztreonam/avibactam (ATM/AVI) is a recently approved combination therapy intended to be used for MBL-producing Enterobacterales and *Stenotrophomonas maltophilia* (16,17). Aztreonam (ATM) is a monobactam that targets PBP3 and is not hydrolyzed by MBLs (5,18,19). Since MBL-expressing bacteria also co-produce SBLs active against ATM, avibactam (AVI) is required to inhibit these enzymes and protect ATM (20). Cefiderocol (FDC) is a siderophore-modified cephalosporin that is refractory to the action of the MBLs VIM-2 and IMP-1 (21–23). Unfortunately, NDM variants hydrolyze FDC (24), and this activity has resulted in treatment failure using this last-resort drug (25,26). The life of β-lactam compounds such as FDC could be prolonged by combination with MBL inhibitors. New boronate-based inhibitors, such as taniborbactam (TAN) and xeruborbactam (XER), were designed to mimicking a tetrahedral transition state and binding the active site of MBLs (27–31). These developments resulted in two possible combination therapies, in different stages of clinical trials: cefepime/taniborbactam (FEP/TAN, NCT06168734) and cefiderocol/xeruborbactam (FDC/XER, NCT07104162). Finally, the penicillin mecillinam (MEC) (targeting PBP2) has been successfully used to treat urinary tract infections of Enterobacterales producing NDM-1 (32,33). Despite NDM variants are generally active against penicillins, this approach represents an alternative therapy, particularly favored by the recent development of the oral prodrug pivmecillinam (34).

The introduction of these new therapeutic options for MBL producers is expected to impose novel evolutionary pressure on clinical strains. We therefore decided to explore the evolutionary potential of NDM toward these clinical challenges to anticipate possible resistance scenarios to last-resort treatments and compare the selected mutations with those present in the already known NDM alleles.

Natural NDM variants have been selected based on their ability to confer resistance in zinc-limiting environments, i.e., upon restriction of the metal cofactor essential for the activity and stability of MBLs in the periplasm (35,36). There are 94 reported NDM variants, which present similar resistance phenotypes in zinc-rich media (5,37). Instead, most NDM variants are better suited than NDM-1 to resist zinc starvation. Zinc binding to MBLs in the periplasm is challenged by the host nutritional immunity response that elicits metal deprivation at the infection site (38,39). Therefore, zinc limitation is considered the main driving force of selection of NDM variants.

Directed evolution is a powerful tool to explore whether these therapeutic pressures can be escaped by accessible NDM substitutions (40–44). To address the evolutionary landscape of NDM toward new therapeutic options, we generated a large library of *bla*_NDM_ variants and subjected it to different selection conditions. Here, we show that NDM-1 has a therapy-dependent adaptive potential. Zinc limitation selected variants that recapitulate natural evolution, validating the robustness and the predictive value of this library. MEC selected a narrow adaptation route centered on substitutions N220Y and M67L that optimize substrate accommodation into the active site. FEP/TAN selected several inhibitor-escape pathways centered on residues E152 and (secondarily) K211, that impair binding of TAN. In contrast, FDC/XER and ATM/AVI imposed stringent evolutionary barriers to NDM evolution, and we could not identify any NDM variant able to provide resistance to these β-lactam/β-lactamase inhibitor combinations. These results define a functional evolutionary map for NDM and provide a framework for anticipating which therapeutic strategies are most vulnerable to escape mechanisms restricted to enzyme optimization.

## Results

### Construction and validation of the library

To comprehensively map the mutational landscape of NDM-1 and explore its adaptive potential under different evolutionary pressures, we aimed to construct a highly diverse randomized library. The mutagenesis protocol was calibrated to target a moderate-to-high mutation rate of approximately 5 mutations per gene, a range expected to allow the exploration of epistatic interactions between distal residues while preserving baseline structural functionality. Accordingly, we generated a randomized library by error-prone PCR (EP-PCR) (45) on the mature *bla*_NDM-1_ coding region and cloned the products into the pMBLe-Δ1 plasmid constructed *ad hoc*. This medium-copy plasmid enables expression of NDM-1 induced by IPTG at levels comparable to those observed in clinical isolates (46). NDM mutants were cloned with the native signal peptide, that targets NDM to the periplasm and includes a lipobox sequence that results in a membrane-anchored protein. The library, transformed in *E. coli*, yielded approx. 20,000 colonies per plate in a total of 65 transformation plates, corresponding to an estimate of 1.3 × 10⁶ individual clones. Sequencing twenty selected clones revealed an average of 5.6 mutations in the *bla*_NDM_ gene, that were randomly distributed throughout the amplified fragment (**Fig. S1**), with a mutation bias similar to that observed for Mn(II) enhanced EP-PCR (45).

Based on the mutation rates observed in the sequenced clones from the library, the PEDEL-AA server predicted that, out of the 4,788 possible point mutations (19 substitutions for each amino acid position), 3,319 would be represented within the library, either individually or in combination with additional mutations, corresponding to 69.3% of all possible point mutations in the mature enzyme. Furthermore, the analysis predicted the presence of 1,681 clones differing from NDM-1 by distinct single point mutations, as well as a total of 8.66 × 10^5^ unique clones in the library. Therefore, this library is expected to be highly diverse and representative of the mutational space available for NDM-1 adaptation under the selection conditions.

*E. coli* TOP10 cells were used for screening the library as a model for Enterobacterales. Phenotypic selections were performed by plating a number of cells on the order of the estimated library size under each condition and comparing the library with an isogenic wild-type NDM-1 control, to identify clones expressing variants with an improved performance. To decipher the genetic changes responsible for these enhanced phenotypes, 50 colonies were isolated for each screening condition, and their entire *bla*_NDM_ coding region was sequenced. The WT *bla*_NDM_ gene was not found in any of the selections that led to variants with optimized phenotypes. Immunoblot analysis of *E. coli* cells expressing representative variants conferring improved resistance phenotypes was performed to assess whether these phenotypes could be due to increased expression levels. In all tested cases, expression levels were similar to those observed for WT NDM-1, revealing that the MIC values can be attributed to specific protein traits (**Fig. S2**).

### Zinc starvation selects clinical NDM variants

We first tested whether the library was able to reproduce the main driver of NDM evolution: resistance under zinc-limited conditions. The screening involved examining the resistance phenotype against cefotaxime (CTX) in the presence of increasing concentrations of the chelator dipicolinic acid (DPA). On plates supplemented with 500 μM DPA, the library sustained growth at CTX concentrations up to 8 μg/mL CTX, compared to 4 μg/mL for the wild-type control. This difference was higher under more severe metal restriction (750 μM DPA), in which growth of bacteria expressing NDM-1 was completely inhibited at 0.50 μg/mL CTX, whereas the library produced variants able to sustain growth up to 4 μg/mL CTX. The selection was focused on the highest antibiotic concentrations where the library outperformed the wild-type control.

Sequencing of the recovered colonies revealed a strongly consistent and convergent mutational pattern (**Table S1**). M154L was the dominant substitution in the selected clones, either alone or in combination with additional mutations (76.9 %) (**Fig. 1 A, B**). M154L is also the most recurrent substitution in nature, being present in 55 % of the clinical NDM variants (11), and has been associated with improved resistance under zinc limitation (such as in NDM-4, NDM-5 and NDM-7, for example) (35,36). Substitutions at E152 were repeatedly recovered as E152K, E152V, E152A or E152R. Other selected positions included D130, D95, M265, N103 and W271. Substitutions in positions 130 and 152 have been documented in natural NDM variants with frequencies of 14 % and 4 %.

**Fig 1.**
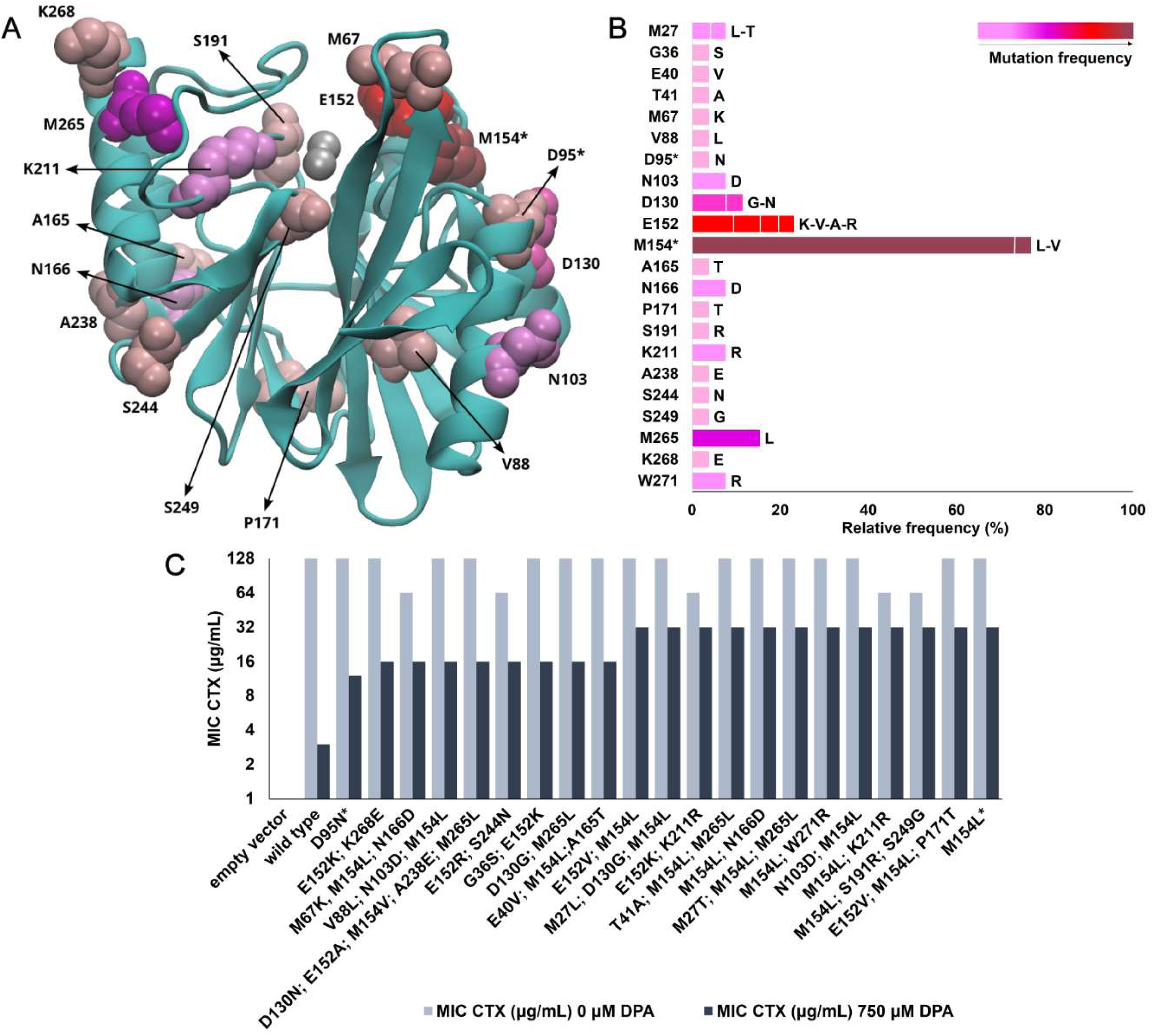
Zinc starvation selects clinical NDM variants. **(A)** Structure of NDM-1 (PDB: 3spu) showing the location of the mutations present in the clones selected with DPA. The protein is shown in cyan. The active-site Zn(II) ions are represented as gray spheres. Mutated residues are represented as spheres of different colors according to the frequency observed for the corresponding substitution as shown in panel B. Positions appearing as a single mutation are marked with an asterisk (*). **(B)** Relative mutation frequency for each amino acid position in NDM-1. Bars indicate the relative frequency (%) of mutations at each position. Observed amino acid substitutions are indicated for each bar: for positions with a single amino acid substitution, the substituted residue is shown at the end of the bar, whereas for positions with multiple alternative amino acids, the bar is subdivided into segments proportional to the frequency of each specific amino acid replacement which are listed at the end of the bar in order of decreasing frequency (or in equal proportions when frequencies match). The color gradient reflects the mutation frequency in the position, ranging from pink (lowest frequency) to dark red/brown (highest frequency). **(C)** MICs of CTX in *E. coli* cells expressing the different NDM variants determined in zinc-replete (0 µM DPA) and zinc-limiting conditions (750 µM DPA). The results correspond to three independent determinations (n = 3). Empty vector refers to pMBLe-Δ1 plasmid without *bla*_NDM_ gene.

MIC determinations confirmed that the selected variants were more resilient to zinc deprivation (**Table S2, Fig. 1C**). In zinc-replete medium (without added DPA), all selected variants exhibited resistance levels equivalent to NDM-1, clustering around a MIC of 128 μg/mL CTX. In contrast, at 750 μM DPA, the resistance phenotype of NDM-1 was severely impaired, with an MIC <1 μg/mL CTX, whereas the selected clones displayed MIC values between 16 and 32 μg/mL CTX.

These findings demonstrate that exposing the mutant library to metal limitation as a selective pressure mirrors the natural evolutionary trends and the evolutionary bottlenecks encountered by pathogens during clinical infection and host immune response. The overlap between laboratory-selected mutants and circulating clinical variants reveals that this library is able to reproduce a central feature of clinical NDM evolution. We therefore expect it to have a powerful predictive potential to anticipate the emergence of future antibiotic-resistant variants.

### Mecillinam selection reveals a narrow evolutionary pathway optimizing substrate binding

The MIC of MEC toward *E. coli* cells expressing WT NDM-1 was 4-8 μg/mL. The library was exposed to increasing concentrations of MEC in plates, showing sustained growth up to 128 μg/mL. The selection was focused on the highest antibiotic concentrations where the library outperformed the wild-type control. Sequencing of the recovered colonies revealed a highly restricted mutational spectrum (**Table S3**). N220Y was dominant, being present in ca. 80% of all analyzed clones in combination with other substitutions distributed all over the protein structure (**Fig. 2 A, B**). The MICs of MEC in *E. coli* cells expressing these variants ranged between 64 and 128 μg/mL, suggesting an important gain-of-function by the acquisition of this substitution (**Table S4, Fig. 2C**). A detailed analysis of the substitutions in the predominant clones shows that only two of them lack N220Y, but share the M67L substitution, with a frequency of 20%. These variants show an MIC of 32 μg/mL. We conclude that enhanced MEC resistance is achieved by one dominant pathway (with a large increase in MIC values), and a second, alternative route, each one with specific substitutions.

**Fig 2.**
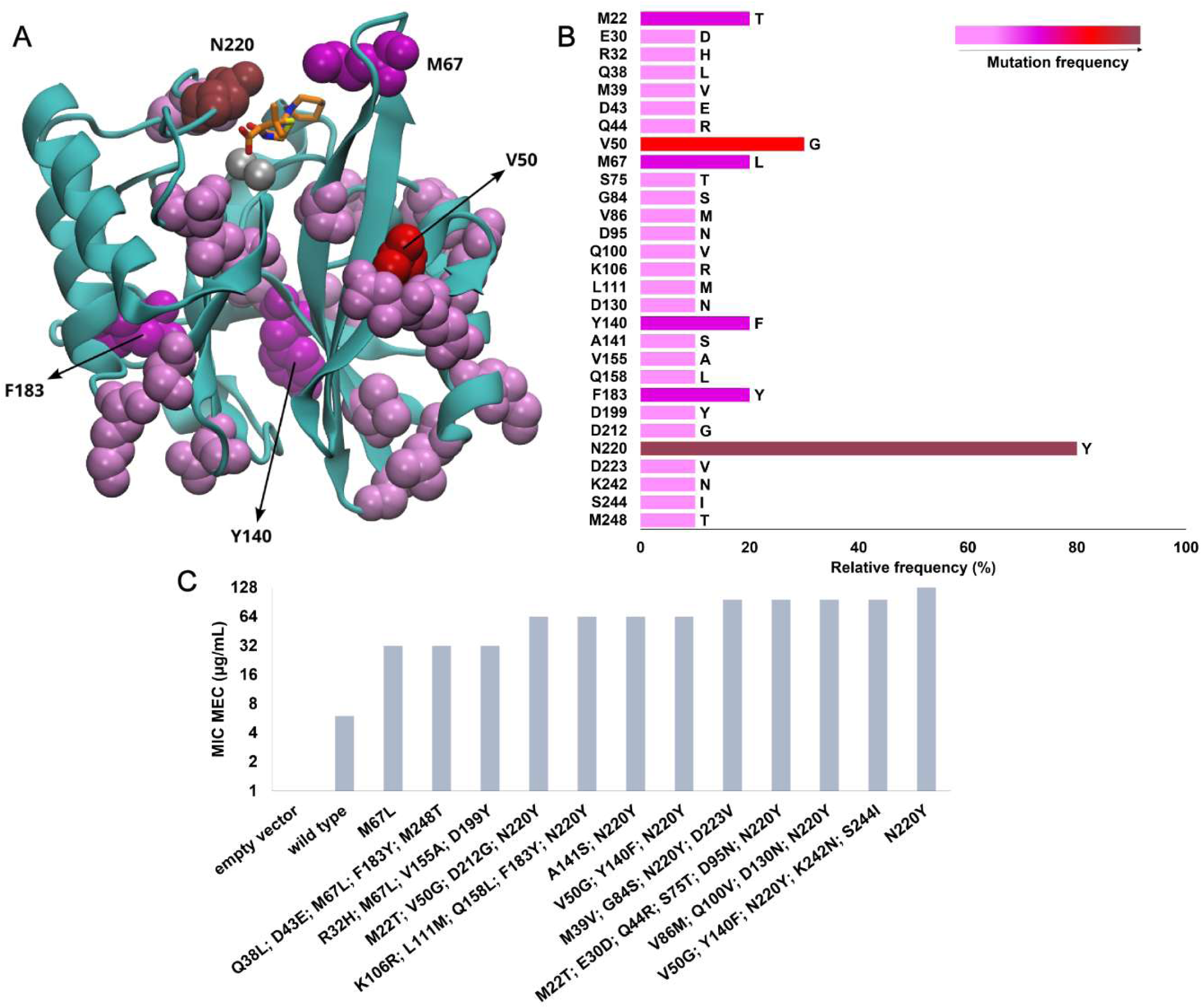
Mecillinam selection reveals a narrow evolutionary pathway. **(A)** Structure of NDM-1 (PDB: 3spu) docked with MEC showing the location of the mutations present in the clones selected. The protein is shown in cyan, while MEC is represented as sticks with carbon atoms in orange, oxygen in red, nitrogen in blue, and sulfur in yellow. The active-site Zn(II) ions are represented as gray spheres. Mutated residues are represented as spheres of different colors according to the frequency observed for the corresponding substitution as shown in panel B. **(B)** Relative mutation frequency for each amino acid position in NDM-1. Bars indicate the relative frequency (%) of mutations at each position. The amino acid substitutions observed are listed at the end of each bar. The color gradient reflects the mutation frequency, ranging from pink (lowest frequency) to dark red/brown (highest frequency). **(C)** MICs of MEC in *E. coli* cells expressing the different NDM variants. The results correspond to three independent determinations (n = 3). Empty vector refers to pMBLe-Δ1 plasmid without *bla*_NDM_ gene.

Reconstruction of the N220Y and M67L single-point mutants confirmed the independent contribution of both substitutions. Expression of NDM-1 conferred a MIC of 4-8 µg/mL MEC. M67L increased the MIC to 32 µg/mL, whereas N220Y alone increased the MIC to 128 µg/mL, fully recapitulating the high-resistance phenotype found in the clones selected in the screening. The additional mutations present in the library isolates do not seem to be essential for the high resistance phenotype, although they may modulate fitness or expression in individual clones.

Computational simulations provide a structural interpretation of the impact of these substitutions in the resistance phenotype. Docking of MEC (**Fig. S3**) results in a larger frequency of productive poses for the N220Y variant compared to wild-type NDM-1 (98% vs. 43%), also correlated with a slightly more favorable binding energy (differing by 0.3 kcal/mol). QM/MM molecular dynamics (MD) simulations on the productive docking pose showed that the L3 loop moved closer to MEC in the N220Y variant, wrapping around the bound substrate and increasing the number of interactions with a hydrophobic patch at the base of loop L3 including residues M67, F70 and V73 (**Fig. 3B**). This rearrangement was not observed in wild-type NDM-1 (**Fig. 3A**). We propose that this conformational change is driven by the interaction of Y220 with the azepane group of MEC, stabilizing the closed active-site arrangement. Binding free-energy calculations show the favorable contribution of residues M67, F70, K211, and N220/Y220 in the N220Y variant compared to NDM-1 (**Fig. S4**), mostly residues from loop L3 and L10.

**Fig 3.**
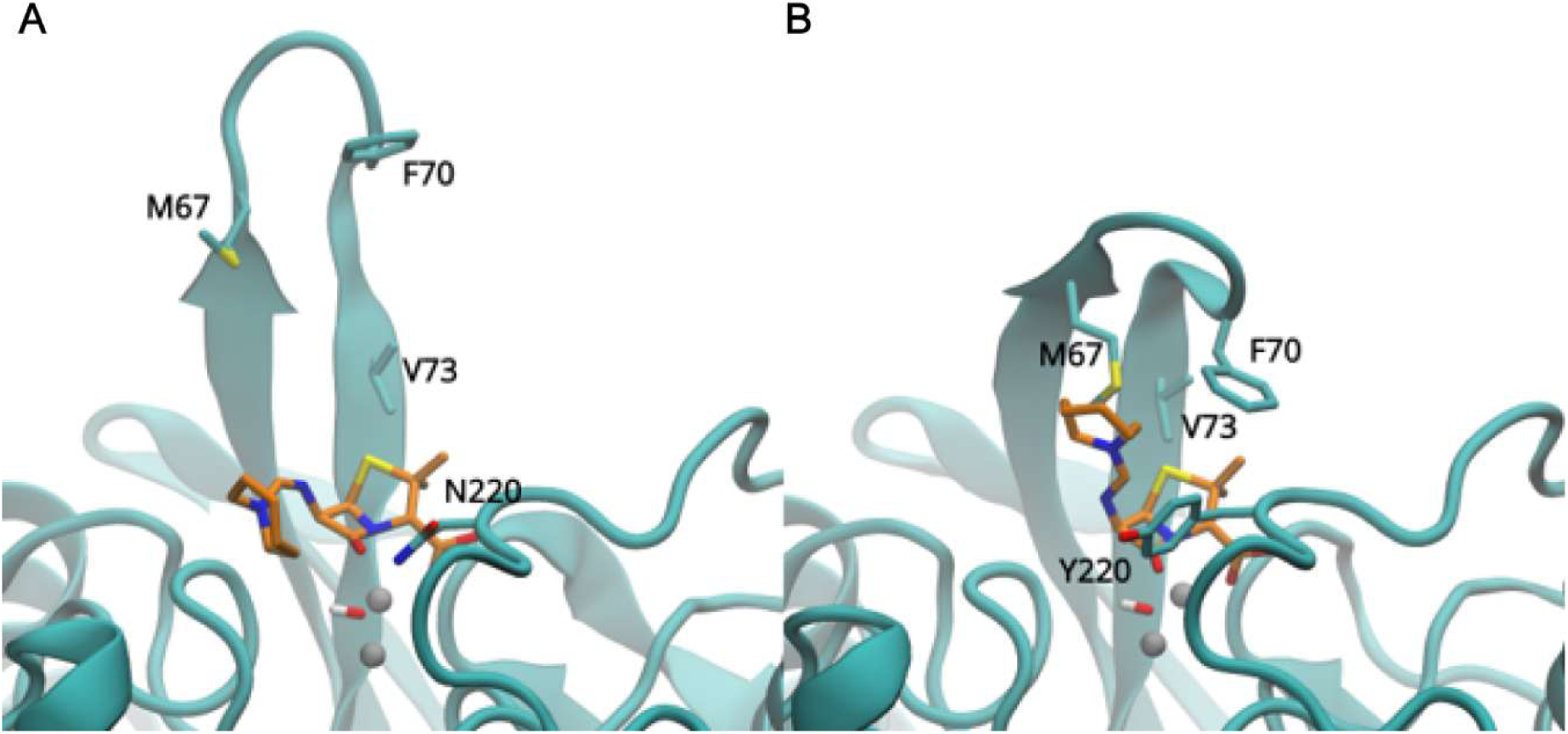
Representative structures obtained by QM/MM MD simulations of mecillinam bound to (A) NDM-1 and (B) the N220Y variant. The proteins are shown in cyan, with selected residues and MEC represented as sticks. Carbon atoms of the residues and MEC are colored in cyan and orange, respectively; oxygen, nitrogen, and sulfur atoms are shown in red, blue, and yellow, and zinc ions as gray spheres.

The M67L substitution in the hydrophobic pocket involves the replacement of a flexible methionine residue by a branched, more hydrophobic leucine. This change can optimize the hydrophobic packing or widen the pocket to better accommodate the bulky amidinopenicillanic acid core of MEC. Hydrophobic substitutions at the base of loop L3 have been already shown to improve the resistance phenotype in clinical variants of the IMP family (47,48). This alternative evolutionary pathway underscores how remote or loop-associated mutations can fine-tune the substrate specificity of NDM-1 without directly disrupting the primary zinc-coordinating residues.

Overall, we conclude that the N220Y substitution enhances MEC recognition by promoting a ligand-induced rearrangement of the L3 loop that strengthens hydrophobic interactions and stabilizes a more productive substrate-binding mode. This provides a molecular explanation for the strong enrichment of N220Y under MEC selection and for the increased MEC MICs observed experimentally and supports the alternative evolutionary path involving M67L.

### Cefepime/taniborbactam selects multiple inhibitor-escape pathways impairing binding of taniborbactam

The phenotypic selection against this combination of a cephalosporin with a boronate inhibitor was performed at a fixed FEP concentration of 1 µg/mL, while varying TAN concentrations from 0 to 64 µg/mL. Under this selective pressure, bacteria expressing NDM-1 grew only up to 0.50 µg/mL TAN, whereas the library yielded resistant variants capable of surviving up to 32 µg/mL TAN, confirming viable evolutionary escape routes against this drug combination.

Sequence of the selected clones revealed a broad mutational landscape scattered across the protein structure, with position E152 as the dominant hotspot (ca. 60%). Substitutions at this residue (including E152G, E152V, and E152K) were recurrently selected, collectively representing most of the isolated substitutions present in resistant population (**Table S5**). The structural mapping and frequencies are presented in **Fig. 4 A, B.** Other few mutations are present with a frequency of ca. 10%. Among them, the most relevant from the biochemical point of view are substitutions in position K211, a residue involved in substrate recognition and also reported to alter binding of TAN (49).

**Fig 4.**
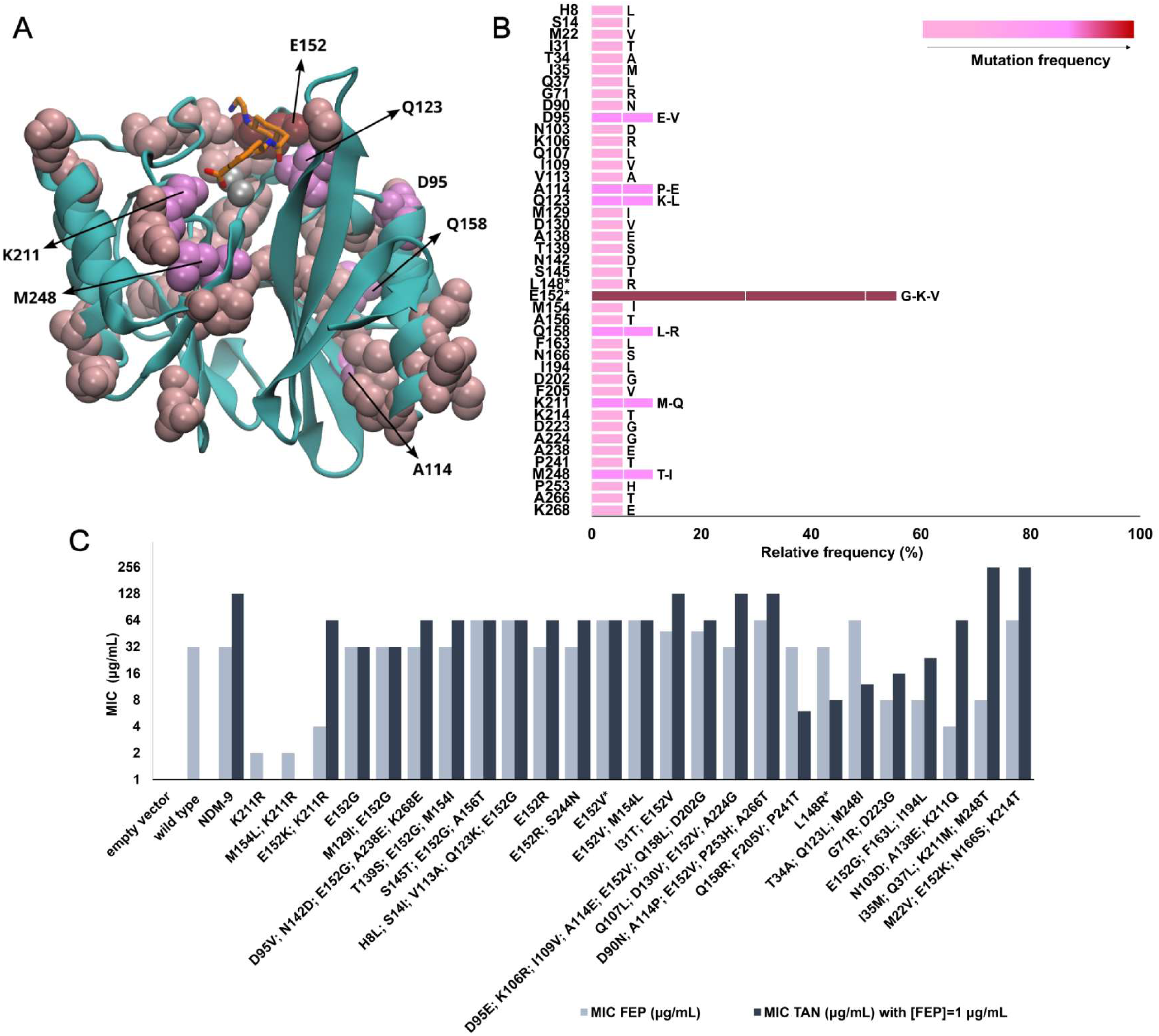
Cefepime/taniborbactam selects multiple inhibitor-escape pathways. **(A)** Structure of NDM-1 (PDB: 3spu) docked with TAN showing the location of the mutations present in the clones selected. The protein is shown in cyan, while TAN is represented as sticks with carbon atoms in orange, oxygen in red, nitrogen in blue, sulfur in yellow, and boron in pink. The active-site Zn (II) ions are represented as gray spheres. Mutated residues are represented as spheres of different colors according to the frequency observed for the corresponding substitution as shown in panel B. Positions appearing as a single mutation are marked with an asterisk (*). **(B)** Relative mutation frequency for each amino acid position in NDM-1. Bars indicate the relative frequency (%) of mutations at each position. Observed amino acid substitutions are indicated for each bar: for positions with a single amino acid substitution, the substituted residue is shown at the end of the bar, whereas for positions with multiple alternative amino acids, the bar is subdivided into segments proportional to the frequency of each specific amino acid replacement which are listed at the end of the bar in order of decreasing frequency (or in equal proportions when frequencies match). The color gradient reflects the mutation frequency in the position, ranging from pink (lowest frequency) to dark red/brown (highest frequency). **(C)** MICs of FEP and MICs of TAN _(FEP= 1 mg/mL)_ in E. coli TOP10 cells. The results correspond to three independent determinations (n = 3). Empty vector refers to pMBLe-Δ1 plasmid without *bla*_NDM_ gene.

To evaluate the independent contribution of the substitutions to the resistance pheno-type, the single-point mutants K211R, E152G (the most frequent among changes in this position) and E152R (also isolated in the DPA selection) (**Table S1**) were constructed. In the wild-type background, TAN restored FEP susceptibility at 1 µg/mL TAN (**Table S6 and Fig. 4C**). In contrast, E152G, E152R, and E152V increased MICs of TAN_(FEP= 1 mg/mL)_ to 32-64 µg/mL, while combinations containing E152K or more complex mutational backgrounds reached higher values. Clones carrying the I35M; Q37L; K211M; M248T and M22V; E152K; N166S; K214T mutation profiles/combinations showed MICs of TAN_(FEP= 1 mg/mL)_ >256 µg/mL, indicating that certain combinations of substitutions in different regions of the protein that do not involve position E152 can elicit strong inhibitor escape. K211R alone had little effect, but K211M and K211Q in selected backgrounds suggest that the positive charge of K211 contributes to inhibitor binding depending on the specific substitution and the mutational context.

The mechanistic interpretation is consistent with chemistry of TAN recognition by MBLs (49–52). In wild-type NDM-1, TAN adopts a productive binding pose that preserves interactions with the zinc-binding site and is stabilized by an electrostatic network involving E152 and K211 (Figure 5). E152 contributes its negatively charged carboxylate group to stabilize the positively charged amino group of TAN, whereas K211 helps orient the TAN carboxylate. Accordingly, wild-type NDM-1 showed the most favorable docking profile, with the most negative binding energy and a productive-pose population of 91% (Figure S5).

**Figure 5.**
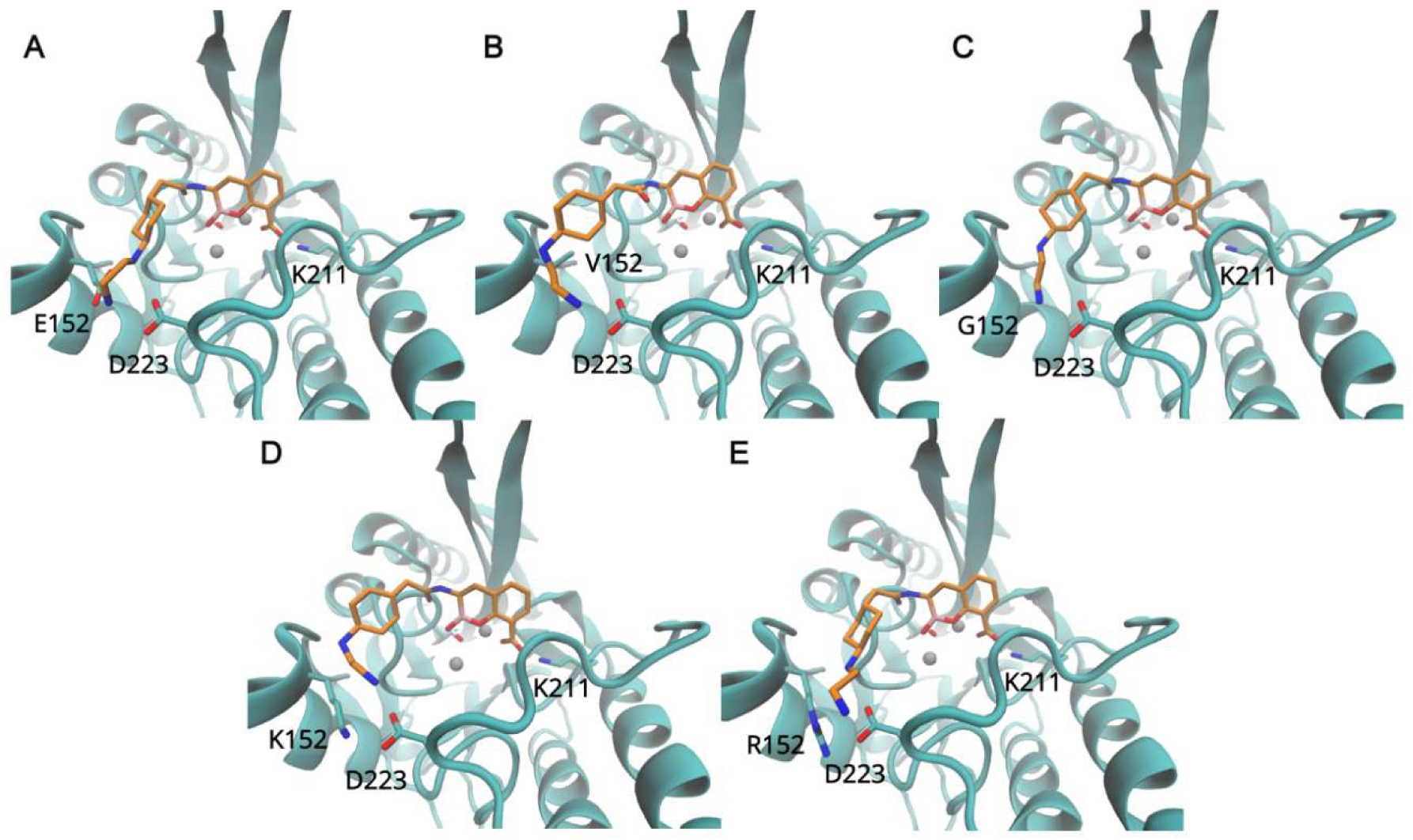
Productive binding poses of taniborbactam in NDM-1 and selected variants. Molecular docking poses of TAN bound to the active site of **(A)** wild-type NDM-1, **(B)** E152V, **(C)** E152G, **(D)** E152K and **(E)** E152R variants. The protein is shown in cyan, while TAN and selected residues are represented as sticks with carbon atoms in orange and cyan, respectively. Oxygen atoms are depicted in red, nitrogen in blue, sulfur in yellow, and boron in pink. Zinc ions are shown as gray spheres.

Substitutions at E152 impaired TAN recognition in a residue-dependent manner. Replacement by glycine or valine removes the favorable negative charge, whereas replacement by lysine additionally introduces a positively charged side chain that can repel the amino group of TAN. Consistent with this model, all E152 substitutions weakened TAN binding energy. E152K showed the strongest effect, combining the least favorable binding energy (ΔΔG = 5.3 kcal/mol) with a reduced productive-pose population of 60%, in agreement with previous observations (39). This result is particularly relevant because E152K defines the clinical variant NDM-9, indicating that the library can recover clinically relevant inhibitor-escape mechanisms while revealing alternative mutational routes at the same position. E152V and E152G retained high productive-pose populations of 97% and 88%, respectively, but showed less favorable binding energies (ΔΔG = 3.3 and 3.6 kcal/mol), confirming impaired TAN binding (**Fig. S5**).

Variants at K211 produced more moderate effects. K211M and K211Q showed less favorable binding energies, with ΔΔG values of 1.3 and 1.5 kcal/mol, respectively, and productive-pose populations of 65% and 71% (**Fig. S5**). These results suggest that loss of the positive charge at position 211 weakens, but does not abolish, productive TAN binding (Figure S6). In contrast, K211R preserved productive binding, consistent with its low resistance profile.

Together, these docking results indicate that variants selected under FEP/TAN pressure impair TAN recognition through localized perturbations of the electrostatic network surrounding the inhibitor-binding site, while preserving the catalytic zinc center.

### Cross-resistance and evolutionary overlap under metal-limiting conditions

Position E152 is a shared evolutionary hotspot developing under both TAN pressure and metal-limitation selection. Therefore, we sought to determine whether the specific amino acid substitutions selected by TAN could also provide a collateral advantage in zinc-depleted environments. To address this, a panel of TAN-selected variants carrying different substitutions at position 152 (E152G, E152K, and E152R) was evaluated against CTX in the presence of 750 µM of DPA (**Fig. 6**).

**Fig 6.**
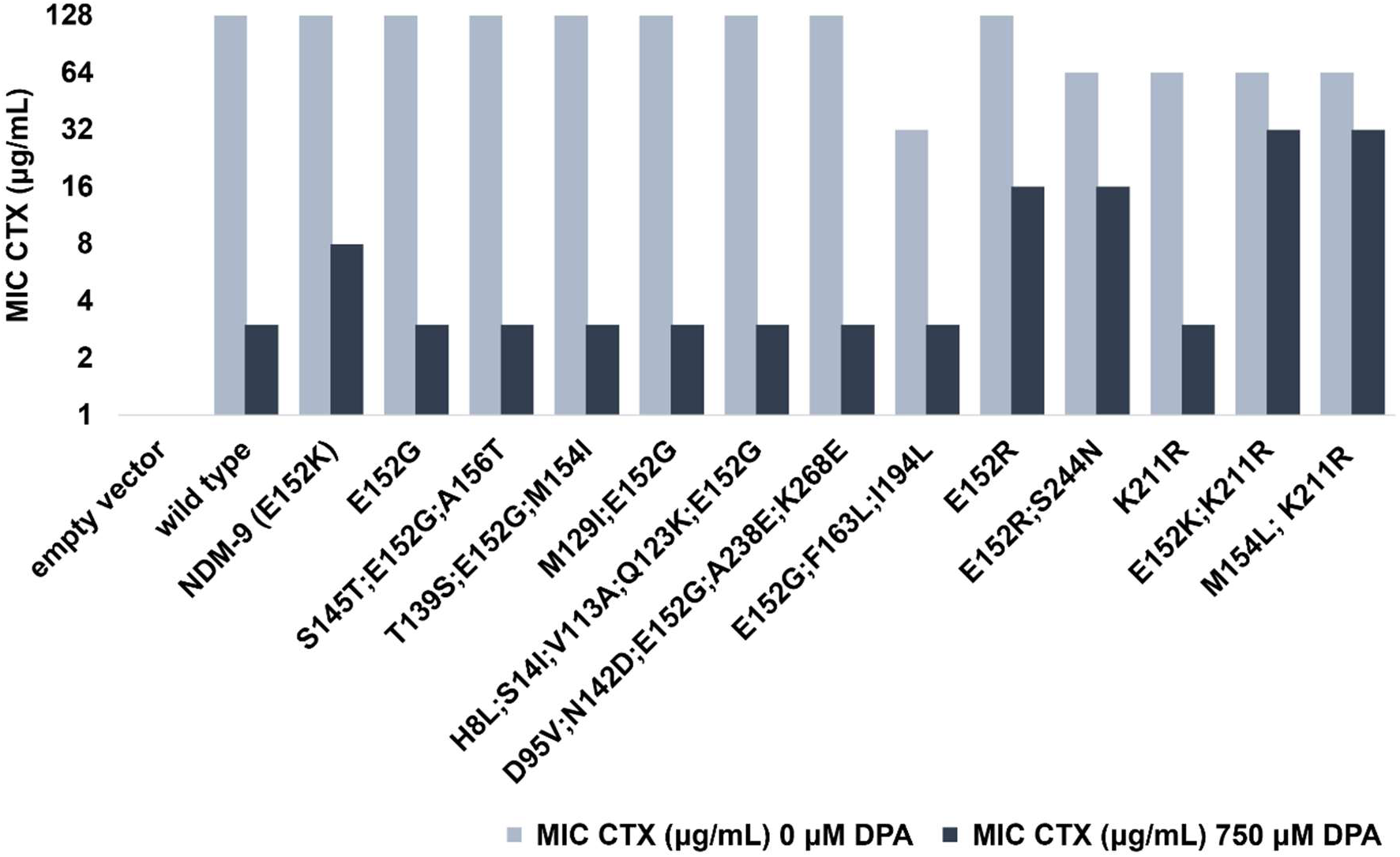
MICs of CTX in *E. coli* cells of a panel of taniborbactam-selected variants carrying different substitutions at position 152 determined in both non-limiting (0 µM DPA) and zinc-limiting conditions (750 µM DPA). The results correspond to three independent determinations (n = 3). Empty vector refers to pMBLe-Δ1 plasmid without *bla*_NDM_ gene.

The most prevalent TAN-selected substitution, E152G, maintained a CTX MIC profile identical to that of wild-type NDM-1 under severe zinc limitation (MIC = 3 µg/mL), both as a single mutant and when accumulated alongside background mutations in various isolated clones. This indicates that while E152G limits inhibition by TAN, it represents a neutral trait toward metal homeostasis, without offering any advantage under low-zinc conditions.

In sharp contrast, substitutions that introduced a positive charge at this position, such as E152K (corresponding to the clinical variant NDM-9) and E152R, significantly enhanced the NDM tolerance to zinc starvation, raising the CTX MIC under 750 µM DPA to 8 µg/mL and 16 µg/mL, respectively. Strikingly, this metal-resistant phenotype was further pronounced in the context of other substitutions: the double-mutant combinations E152K/K211R and M154L/K211R displayed a remarkable synergistic effect, driving the CTX MIC up to 32 µg/mL upon zinc deprivation. This reveals a high potential for adaptation of these mutations while instead, E152G has a reduced competitive advantage. Taken together, these cross-resistance patterns reveal an evolutionary overlap between inhibitor escape and adaptation to zinc restriction, where specific structural pathways selected under TAN pressure can co-select for highly resilient phenotypes capable of overcoming severe nutritional immunity bottlenecks.

### Cefiderocol/xeruborbactam and aztreonam/avibactam impose high evolutionary barriers

Phenotypic screening under the combination of cefiderocol and xeruborbactam (FDC/XER) led to a completely distinct evolutionary outcome. This screening was performed using the Iron-Depleted Cation-Adjusted Mueller-Hinton broth (ID-CAHMB) medium, specific for FDC. Under these screening conditions, *E. coli* cells expressing WT NDM-1 tolerated XER concentrations up to 0.125 µg/mL; however, no clone from the library displaying improved resistance was recovered. This negative selection is informative since the library was large enough to provide adaptive routes under zinc limitation, MEC or FEP/TAN. This result suggests that XER places a strong constraint on the accessible NDM mutational space under these conditions.

The combination of aztreonam with the SBL inhibitor avibactam (ATM/AVI) is the treatment of choice for MBL-producing Enterobacterales. Screening against ATM alone and in combination with AVI produced an equally restrictive result. Neither the wild-type NDM-1 control nor the library generated colonies at ATM concentrations as low as 0.06 µg/mL, and no ATM/AVI-resistant NDM variants were recovered. This result is consistent with the structural limitation that MBLs bind ATM in a nonproductive mode and do not efficiently hydrolyze monobactams.

## Discussion

In this work, we identified the adaptive landscapes of NDM-1 when challenged with the evolutionary pressures of different therapeutic strategies. The widely different results show that NDM evolvability is not a uniform property of the enzyme, but it depends on whether the selective challenge can be solved by local changes in substrate positioning, inhibitor recognition, or tolerance to zinc starvation. Zinc limitation, MEC and FEP/TAN each selected distinct evolutionary outcomes. In contrast, FDC/XER and ATM/AVI did not yield NDM variants able to confer improved resistance compared to NDM-1, indicating a much higher barrier to enzyme-level escape.

The results from the zinc-limitation selection are relevant for two reasons. Mechanistically, it confirms the hypothesis that NDM variants are being selected to overcome metal scarcity. M154L emerged as the most frequent substitution, and E152 appeared as a recurrent secondary hotspot. These results agree with the view that clinical NDM evolution is shaped by the need to preserve function under host-imposed zinc restriction. Methodologically, the recovery of clinically convergent substitutions validates the quality of the library. Indeed, NDM variants can overcome the challenge of zinc restriction by improving two traits in evolution: (1) improving the zinc binding affinity, a feature exclusively provided by substitution M154L (36) or (2) improving the stability of the zinc-deprived enzyme in the periplasm (53), as achieved by substitutions E152K, D130G/N, V88L, among others. Here we isolated variants with mutations that impact both mechanisms, in contrast to a previous direct evolution study in which the most prevalent clinical substitution, M154L, was not isolated (43).

The selection against mecillinam shows how NDM can adapt to improve the resistance phenotype of a substrate that belongs to a family of β-lactam compounds (penicillins) already recognized and cleaved by NDM. This was achieved by incorporating changes in the loops surrounding the active site. The strong enrichment of N220Y (at loop L10) and the phenotype of the reconstructed single mutant support a simple model: substitution at position 220 improves substrate accommodation by closure of the active site loop L3 and favoring hydrophobic interactions between MEC and the enzyme active site. Since N220 is not the dominant route in zinc-starvation or TAN selections, MEC resistance is a therapy-specific adaptation. The secondary adaptive route, involving the M67L mutation, improves the same structural trait favored by N220Y, i.e., hydrophobic recognition in the active site, disclosing a convergent evolution from the biochemical point of view. From a clinical perspective, this finding warns that renewed use of MEC, especially in settings where NDM-producing Enterobacterales circulate, could select uncommon but accessible NDM variants with improved substrate recognition and resistance phenotype.

Taniborbactam produced the most clinically concerning adaptive pattern. The selection converged on E152, a position already implicated in TAN resistance through NDM-9 (52,54), but it also revealed that loss or reversal of the electrostatic contribution involving E152 can be achieved through several substitutions. This enforces the notion that TAN escape is not limited to one clinical variant, since multiple nucleotide routes can erode the E152-mediated interaction network. In addition, non-conservative changes in position K211 can further impair inhibitor recognition. A comprehensive randomized single codon mutations study revealed that the essentiality of K211 is strongly substrate ion dependent, in agreement with our findings with TAN (55). This work also reveals other combinations of mutations that can effectively reduce the inhibitory impact of TAN by less obvious mechanisms. In summary, these results suggest that the therapeutic use of FEP/TAN (if approved) should be accompanied by surveillance for E152 substitutions, careful susceptibility testing, and awareness that inhibitor resistance may arise without a major loss of FEP resistance conferred by NDM.

The contrast with cefiderocol/xeruborbactam (which is also a combination of a cephalosporin and a boronate-based inhibitor) is striking. Under our screening conditions, we could not find any variant outperforming wild-type NDM-1. Directed evolution of NDM-1 towards FDC led to the identification of mutations that did not improve the resistance phenotype (56). Here we show that combination of this siderophore-modified cephalosporin with XER constrains the accessible NDM mutational space more effectively than TAN. An in vitro obtained S262G variant of NDM-1 was shown to be resistant to the combination meropenem-XER (57). The absence of this mutation in the current screening can be accounted for a possible higher challenge of the FDC/XER couple compared to meropenem/XER, but also to the possibility that this substitution can lead to an impaired expression or destabilization of NDM that may not be able to provide resistance to these conditions. Our results support further development of XER-containing combinations for MBL-producing organisms. However, this observation should be framed carefully. FDC efficacy can be affected by iron transport, permeability, PBP3 changes, β-lactamase expression level, and species-specific physiology. Therefore, the absence of NDM enzyme escape routes in this work cannot be extrapolated to infer absence of clinical resistance. The minimalistic structure of XER compared to TAN leaves few contact points of this inhibitor with active site residues, possibly limiting the evolutionary accessible escape routes that can provide an impaired XER binding while preserving the enzyme stability and hydrolytic capability against FDC.

Aztreonam/avibactam appears highly robust from the perspective of NDM evolution. The failure to recover improved NDM variants is consistent with the long-standing observation that MBLs do not efficiently hydrolyze monobactams. AVI protects ATM against co-produced serine β-lactamases, converting a biochemical blind spot of MBLs into a therapeutic strategy. Recent works have reported the existence of MBL-related enzymes with ability to hydrolyze aztreonam (58,59). However, these enzymes are evolutionarily distant to B1 MBLs and show impaired activity against cephalosporins and carbapenems. This reveals that the possible acquisition of activity against ATM of NDM can take place at a high cost, involving an intense active site remodeling that may pose a large evolutionary barrier. We conclude that is unlikely that NDM can easily evolve activity against ATM. These results suggest that the main clinical threat for this therapy might not be linked to NDM evolution, but to PBP3 insertions, permeability changes, efflux or altered expression of co-produced serine β-lactamases.

Overall, our data show a hierarchy of enzyme-related evolutionary risks in different therapies. Zinc limitation is a powerful background pressure that selects NDM variants during infection. FEP/TAN shows the clearest NDM-mediated escape potential. MEC can be better hydrolyzed through a narrow but potent substrate-adaptation route. In contrast, ATM/AVI and FDC/XER impose high barriers to NDM-centered evolution. These findings support therapeutic strategies that prioritize combinations forcing resistance to move outside the accessible NDM mutational landscape, while integrating WGS strategies cell and clinical surveillance for non-MBL mechanisms. Finally, the finding of this broadly divergent adaptive landscape further highlights the need for different therapeutic choices to treat MBL producers

## Author contributions

FDR, LJG and AJV designed the research; FDR performed the selections and analysis for all conditions, MICs assays, immunodetection analysis and construction of mutants; GB and GG constructed the library; SID and DMM designed, performed, analyzed the computational experiments and wrote the corresponding text sections; FDR, LJG and AJV analyzed and discussed the data; FDR, LJG and AJV wrote the paper; all authors discussed ideas, reviewed and edited the manuscript.

## Acknowledgments

This work was supported by grants PICT-2016-1657 and PICT 2020-03769 from Agencia I+D+i to AJV and DMM, PICT 2020-1923 to LJG, PID 80020220700052UR from UNR to DMM, ASaCTeI (PEICID 2023-191) and the REPARA network grant (Redes Federales de Alto Impacto, Subsecretaría de Ciencia y Tecnología de la Nación) to AJV. Funds and/or facilities were provided by the Cleveland Department of Veterans Affairs, award number 1I01BX001974 to RAB., from the Biomedical Laboratory Research & Development Service of the VA Office of Research and Development and the Geriatric Research Education and Clinical Center VISN 10 to RAB. AJV, LJG, SID and DMM are staff members of CONICET. We are grateful to Liliana Rojas (IBR-CONICET) for her excellent technical assistance. The content is solely the responsibility of the authors and does not necessarily represent the official views of the NIH or the Department of Veterans Affairs.

## Materials and Methods

### Bacterial strains and growth conditions

*Escherichia coli* TOP10 (Invitrogen) was employed for the generation and assembly of the library. Cells were routinely grown aerobically at 37°C in Luria-Bertani (LB) broth (Difco) or on LB agar plates, unless otherwise specified. When necessary, gentamicin (Sigma-Aldrich) was added at a final concentration of 20 μg/mL to maintain plasmid selection.

For all susceptibility assays involving cefiderocol, Iron-Depleted Cation-Adjusted Mueller-Hinton Broth (IDCAMHB) was employed prepared according to the Clinical Laboratory and Standards Institute (CLSI) protocol (60). To ensure the correct preparation and quality control of the medium, standard reference strains were concurrently evaluated. The panel included *Acinetobacter baumannii* M27858 CDC #0083 (NDM-1; PER-7; OXA-23; OXA-69; MIC > 32 μg/mL, resistant), *Pseudomonas aeruginosa* M27875 CDC #0064 (NDM-1; OXA-50; PAO; OXA-56; MIC = 2 μg/mL, susceptible) and *Escherichia coli* ATCC 25922 (MIC range of 0.06–0.5 μg/mL).

### Plasmid construction

The plasmid pMBLe-NDM-1-ST was used as the starting construct. This plasmid contains the gene encoding the NDM-1 enzyme flanked by *Nde*I and *Hind*III restriction sites and allows expression of the enzyme fused to a C-terminal Strep-tag II, under the control of an IPTG-inducible pTac promoter. This allows uniform quantification by immunoblotting without affecting the protein localization or the resistance phenotype, and regardless of the mutations introduced (13).

To facilitate the study of the gene region encoding mature NDM, we introduced restriction sites *Pst*I and *Sma*I flanking the lipobox near the end of the enzyme signal peptide. These recognition sites were incorporated through silent mutations via the “Vector amplification mutagenesis” technique described in the next section, generating the modified vector designated pMBLe-NDM-1-ST PS. To improve transformation efficiency, a minimized version of the vector designated pMBLe-Δ1-NDM-1-ST PS, was engineered. The primer pair B1/B2 (**Table S7**) was designed to remove a dispensable 2195 bp fragment located between the origin of replication (*pBBR1 Rep/OriV*) and the *ptac* promoter, including the *lacI* gene. Following amplification, 5′ phosphate groups were incorporated into the PCR products using polynucleotide kinase. The linearized fragments were circularized by blunt-end ligation, yielding a new vector version of 4952 bp.

### Vector amplification mutagenesis

Site-directed mutations were introduced using a vector amplification mutagenesis protocol. Briefly, a pair of complementary oligonucleotides containing the desired mutation were designed to anneal to identical positions on opposite strands of the template plasmid. Extension was performed via PCR utilizing a highly processive, high-fidelity polymerase. The polymerase synthesized the complementary strands by completely copying the circular plasmid template without displacing the mutagenic primers. The resulting unmethylated strands then annealed to form a nicked plasmid. Following the amplification reaction, the parental methylated DNA and hemimethylated hybrid molecules were digested using the *Dpn*I restriction endonuclease. This step enriched the mixture for the unmethylated, nicked mutagenized plasmid product. The digestion products were subsequently transformed into *E. coli* TOP10 competent cells, where the nicked plasmids were repaired by host ligases and replicated.

Single-point mutant variants (M67L, E152G, E152R, K211R and N220Y) were constructed independently to isolate and evaluate the specific biochemical contributions of key hot-spot mutations using the corresponding mutagenic primers and the pMBLe-Δ1-NDM-1-ST PS plasmid. Constructs were verified by PCR using plasmid-specific primers that anneal outside the multiple cloning site (MCS) (**Table S7**).

### Error-Prone PCR (EP-PCR) and Template Optimization

Random mutagenesis of the *bla*NDM-1 gene was performed using an Error-Prone PCR protocol (EP-PCR).(61) The protocol was calibrated to achieve 10 total duplications using a low initial template concentration. In addition, the frequency of each mutation type was quantified (**Table S8**), grouping each type of point substitution together with its complementary change. Prior to mutagenesis, a standard PCR reaction was performed to amplify the *bla*NDM-1 gene from the pMBLe-Δ1-NDM-1-ST PS plasmid. The purified product was then subjected to the optimized EP-PCR conditions.

### Library Construction

For library assembly, both the mutagenized EP-PCR amplicons and the pMBLe-Δ1-NDM-1-ST PS vector were digested using the *Pst*I and *Hind*III enzyme pair. This cloning strategy allows the subcloning of the mature portion of the NDM-1 gene directly into the expression framework. 65 individual aliquots of chemically competent *E. coli* TOP10 cells prepared using the commercial Mix&Go kit (Zymo Research) were transformed and each aliquot was plated onto an individual agar plate.

To evaluate library breadth, the bioinformatic tool PEDEL-AA (available at https://guinevere.otago.ac.nz/STATS/pedel-AA.php) was employed.

### Phenotypic Selection and Minimum inhibitory concentration (MIC) determinations

Phenotypic screening of the library was performed under distinct selective pressures. Mutant selection was performed on LB agar plates by plating approximately 5×10^6^ cells under each condition tested, corresponding to a number on the order of the estimated library size to maximize the probability of capturing all relevant variants. To determine the optimal selection conditions, identical numbers of cells from the library and *E. coli* TOP10 pMBLe-Δ1-NDM-1-ST PS were plated in parallel. This allowed a direct comparison between the growth supported by wild-type NDM-1 and that supported by potential mutant variants.

To simulate physiological zinc-limiting conditions, selection assays were conducted on LB agar plates supplemented with dipicolinic acid (DPA) at concentrations of 500 μM and 750 μM, across a Cefotaxime (CTX) concentration gradient (0–128 μg/mL for 500 μM DPA; 0–16 μg/mL for 750 μM DPA). For novel therapeutic escape profiling, screenings were conducted using Mecillinam (MEC) or the Cefepime/Taniborbactam (FEP/TAN), Cefiderocol/Xeruborbactam (FDC/XER) and Aztreonam/Avibactam (ATM/AVI) combinations (maintaining a fixed FEP concentration of 1 μg/mL while varying TAN from 0 to 64 μg/mL; a fixed FDC concentration of 0.125 μg/mL while varying XER from 0 to 0.5 μg/mL; and a fixed AVI concentration of 1 μg/mL while varying ATM from 0 to 1 μg/mL). Surviving individual colonies capable of growing at concentrations that completely inhibited wild-type control were isolated and sequenced.

Minimum inhibitory concentrations (MIC) of both selected library clones and isolated single-point mutants were determined using broth microdilution method using LB or ID-CAMHB, according to the CLSI protocol (REF). The final concentration of *E. coli* TOP10 cells was adjusted to 5 x 10^5^ CFU/ml and to induce the expression of *bla*_NDM-1_ or the variants from the pMBLe Δ1 plasmid, the bacterial inoculum was prepared with 100 µM of isopropyl β-d-1-thiogalactopyranoside (IPTG). Plates were incubated at 37ᵒC for 18 h. Results shown are the mode of three independent experiments.

### Molecular Docking

Molecular docking simulations were performed using Autodock-GPU (62). The grid maps were set to 70 × 70 × 70 points with a grid spacing of 0.375 Å centered on the oxygen atom of the hydroxyl ion in the catalytic site. For each docking calculation, 100 different docking runs were performed, which were clustered using a RMSD cut-off of 2 Å. Between all the docking poses produced by the docking simulation, the productive pose of each ligand was selected, defined as the one in which there is a distance of less than 3.0 Å between the atoms involved in the hydrolysis reaction at the active site (the carbon atom of the carbonyl group of the beta-lactamic ring in the case of MEC or the boron atom in the case of TAN, and the oxygen atom of the hydroxyl group in the active site).

The structure for NDM-1 was taken from PDB ID 3SPU (63) and all mutants were modelled using the AMBER24 package (64), first by manually mutating the residue with TLeap and then performing a geometry optimization with Sander. The optimization was done with implicit solvent using the Generalized Born method, with all atoms except the ones that belong to the Y220 residue frozen in place by a force constraint of 1000 kcal/mol.

The structures of the ligands were taken from PubChem and processed by optimization of geometry and calculation of RESP charges using Gaussian 16 (65) at B3LYP/6-31G** level. The ligand was then processed using Autodock Tools to prepare it for docking.

### QM/MM Molecular Dynamics Simulations

Hybrid Quantum Mechanics/Molecular Mechanics (QM/MM) molecular dynamics simulations were performed using the PMEMD software from the AMBER24 package. The initial coordinates were taken from the productive pose of MEC on each protein and submerged in a periodic truncated octahedral box of molecules. The QM subsystem consisted of the ligand, the hydroxyl group, the Zn^2+^ ions, and the sidechains of the residues coordinated to those ions (H120, H122, D124, H189, C208, H250), and its calculations were done using the semiempiric DFTB3 method (66) The rest of the system was treated as MM, with ff14SB parameters (67) for the protein and TIP3P for the water molecules. The thermostat and barostat used were Langevin and Monte Carlo, respectively.

The simulation protocol started with 10000 steps of energy minimization, followed by 10 ps of thermalization from 0 to 300 K of temperature at constant volume and 20 ps of equilibration at constant temperature and pressure, finalizing in 1 ns of production at constant temperature and pressure.

### Free binding energy calculations

Binding energy calculations between the proteins and the ligands were performed using the MMGBSA program in AMBER24. These calculations are performed using MM parametrization of the system. This aspect limits the capacity of studying the interactions between the ligand and the Zn^2+^ ions, which involve coordination through the d orbitals of the ions and would require more complex QM calculations, but in this case it is used to characterize the interactions between the ligand and the sidechains of residues in loops L3 and L10, which makes this method suitable. For the MM parameters of the ligand, antechamber was used with the GAFF2 parameters, together with the RESP charges from the QM optimizations from the docking studies, and the parameters of the active site residues and ZN^2+^ ions were taken from bibliography (68).

### Protein immunodetection

Overnight cultures were diluted 1:100 in fresh LB medium and grown at 37°C with shaking until OD_600_ of 0.4, after which IPTG 100 µM final was added. After 2 hours of induction at 37°C and shaking, cell lysates were resolved by SDS-PAGE and transferred to membranes for Western blotting using Strep-Tag® II monoclonal antibodies (Sigma-Aldrich) at a 1:1000 dilution (from a 200 μg/mL stock) followed by incubation with an anti-mouse secondary antibody. Immunodetection of GroEL was used as a loading control.

